# AltORF-encoded microprotein recruits primary-ORF-encoded dehydrogenase to cell poles upon heat shock

**DOI:** 10.64898/2026.10.22.760973

**Authors:** Janka J. Schmidt, Félix Ramos-León, Aixia Zhang, Rilee Zeinert, Svetlana A. Shabalina, Gisela Storz

**Affiliations:** Division of Molecular and Cellular Biology, Eunice Kennedy Shriver National Institute of Child Health and Human Development, Bethesda, MD 20892, USA; Laboratory of Molecular Biology, National Cancer Institute, Bethesda, MD 20892, USA; Computational Biology Branch, Division of Intramural Research, National Library of Medicine, National Institutes of Health, Bethesda, MD 20894, USA

**Author notes:** To whom correspondence should be addressed. Gisela Storz,. The authors declare no competing interest.

**Keywords:** intragenic ORF, RNA thermometer, pentose phosphate pathway, NADPH

## Abstract

Alternative open reading frames (altORFs) initiating from in-frame or out-of-frame start sites within annotated protein-coding genes represent a largely unexplored source of protein-coding potential and remain poorly understood in terms of regulation, function, and impact on physiology. Here we investigate *Escherichia coli* GndA, encoded by an out-of-frame altORF within *gnd*, which encodes 6-phosphogluconate dehydrogenase. We observe that the GndA protein is induced by heat shock, consistent with the initial identification in a quantitative membrane proteomics screen, but is unstable, while the levels of the stable Gnd dehydrogenase are unchanged. Using translational fusion assays and in vivo RNA structure probing, we show that an internal complementary sequence upstream of *gndA* acts as a thermo-responsive secondary RNA structure controlling GndA translation. Mutagenesis of either *gnd* or *gndA* reduce heat shock survival of *E. coli* indicating both gene products are important for resistance. GndA fractionates with membranes consistent with in silico predictions that the protein forms an amphipathic α-helix. In contrast, Gnd is predominantly cytoplasmic but shows GndA-dependent relocalization to cell poles upon heat shock. Together, our results demonstrate that an altORF-encoded microprotein is required for stress-dependent relocalization of its primary ORF partner, revealing a functional interplay between proteins encoded by overlapping ORFs in bacterial stress adaptation.

**Significance:** Alternative translation from internal translation initiation sites enables bacteria to expand their proteomic versatility without increasing genome size. Here we show that a single *Escherichia coli* transcript encoding the well-characterized dehydrogenase Gnd also produces an additional microprotein from an internal alternative open reading frame (altORF), whose translation is controlled by an RNA thermometer-like structure. We further demonstrate that this heat shock-induced microprotein promotes spatial relocalization of its primary ORF-encoded partner during aerobic heat stress. Together our findings highlight how alternative translation can expand the cellular response to stress.

## Introduction

Conventional bacterial gene annotation assumes a one-gene-one-protein paradigm, in which translation from the primary translation initiation site (pTiS) produces the canonical protein. However, bacteria can expand their coding potential without increasing genome size through translation of alternative open reading frames (altORFs) nested within canonical genes (reviewed in (1)). One mechanism of altORF translation involves ribosome frameshifting (reviewed in (2, 3)), which shifts the ribosome reading frame to produce elongated or truncated protein isoforms. Another mechanism for altORF translation is initiation from an internal translation initiation site (iTiS). The iTiS can be in-frame, producing N-terminally truncated isoforms that share the C-terminal sequence with the canonical protein, or out-of-frame, producing proteins with distinct sequences and functions (reviewed in (1)).

At present, only a limited number of altORF-encoded proteins have been identified, but these are found within primary genes that encode proteins with diverse roles in virulence for the *Coxiella burnetii* Mip macrophage infectivity potentiator (4), antibiotic synthesis for *Streptomyces venezuelae* PikA (5), protein quality control for *Escherichia coli* and *Synechococcus* ClpB protein disaggregases (6, 7), chemotaxis for *E. coli* CheA (8), and phage defense for *E. coli* DNA endonucleases Rpn (9), among others. Most of these examples were uncovered incidentally during protein purification or functional analyses, suggesting relative high abundance and/or significant roles. We suggest that alternative translation likely is not a rare anomaly but rather a widespread strategy with an as-yet unknown contributions to bacterial fitness.

Not many altORF-encoded proteins have been detected by global protein identification, likely due to inherent challenges in their discovery such as computational identification of ORF-internal translation initiation sites along with the small size of many of the altORF-encoded proteins. One exception is GndA, a microprotein, encoded in the +1 reading frame with respect to the primary *gnd* ORF. Initially, GndA was identified in a quantitative membrane proteomics study conducted under heat shock conditions (10). Translation of GndA was independently verified by ribosome profiling (11). The primary protein encoded by *gnd* is a 6-phosphogluconate dehydrogenase, a well-characterized enzyme in the pentose phosphate pathway that catalyzes the oxidative decarboxylation of 6-phosphogluconate to ribulose-5-phosphate and CO₂, while reducing NADP⁺ to NADPH (12, 13). Regulation of *gnd* expression was found to be growth rate dependent and proposed to be at a post-transcriptional level more than three decades ago (14). The early studies suggested that an internal complementary sequence (ICS) corresponding to *gnd* codons 67–78, which is complementary to the 5’UTR including the ribosome binding site (RBS) of *gnd*, facilitates the formation of a mRNA secondary structure that could sequester the *gnd* RBS to inhibit translation initiation (15, 16). Given that the primary ORF is mechanistically and biochemically well characterized, *gnd* provides an ideal model for dissecting the regulation and role of alternative translation within a defined genetic and metabolic context.

Using multiple reporter fusions in combination with structure probing, we show that GndA translation is regulated by a temperature-responsive secondary mRNA structure (RNA thermometer) encompassing the previously described ICS (15, 16) within the *gnd* coding sequence. Upon heat shock, the structure selectively permits translation of GndA, highlighting independent translation control from a single mRNA. Deletion of the whole locus as well as selective loss of either Gnd or GndA impairs heat shock survival, suggesting functional interplay between the primary protein Gnd and its altORF-encoded partner GndA. The small protein, predicted to be an amphipathic helix, was found to be associated with the membrane and to recruit the Gnd dehydrogenase to the cell poles specifically upon heat shock. Our findings establish GndA as a model for conditional internal translation in bacteria and document an altORF role in stress adaptation.

## Results

### AltORF-encoded GndA is conditionally translated during aerobic heat shock

GndA was initially identified in a membrane proteomics study (10). Independently, ribosome profiling (11, 17) revealed initiation-associated ribosome occupancy at the *gnd* start codon and at a second internal start codon (Fig. S1A), corroborating translation initiation from an internal region of *gnd*. A putative Shine-Dalgarno-like sequence (5′-AGGAGA) and possible UUG and AUG start codons are near the internal ribosome profiling signal (Fig. 1A). These sequences are followed by an in-frame stop codon (TGA) located 105 bp downstream of the AUG start codon. Translation of the ORF starting with the AUG would lead to a 36-aa protein, referred to as GndA here, in the +1-frame relative to the Gnd coding sequence, (Fig. 1A).

**Fig. 1.**
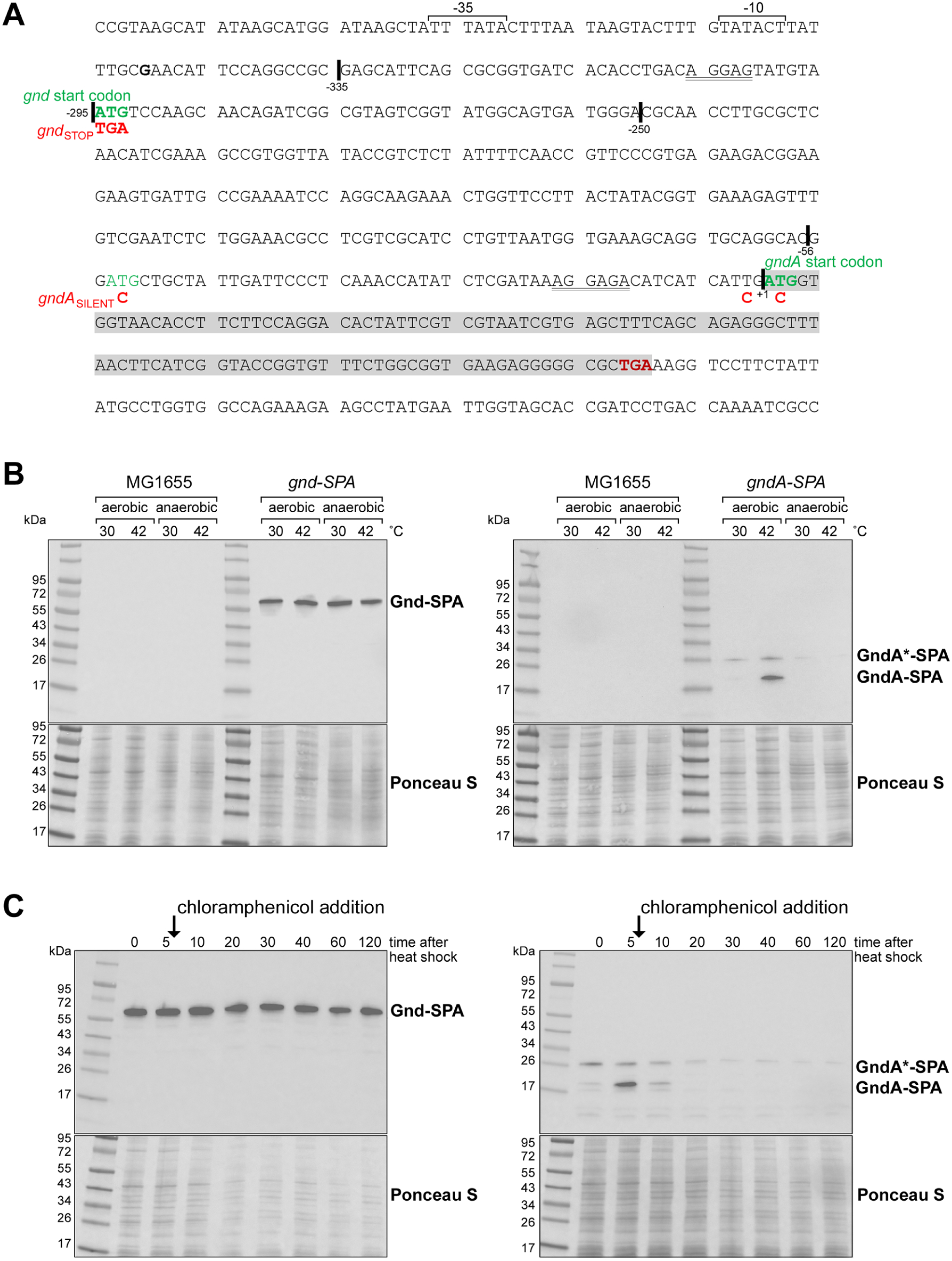
GndA translation is induced by aerobic heat shock. **(*A*)** Coding sequence of *gndA* and 300 nt upstream (with −35 and −10 promoter sequences underlined and +1 nucleotide in bold). Start codon for *gnd* is in green font. Coding sequence for 36-aa *gndA* is highlighted in gray and corresponding AUG start (based on ribosome profiling) and stop codon are in green and red font, respectively. A possible upstream AUG start codon is also in green font. Predicted Shine-Dalgarno sequences are indicated with double underlines. Sequences of *gnd*_STOP_ and *gndA*_SILENT_ mutants are shown. Black lines indicate 5’ ends of constructs assayed in **Fig. S1D**. **(*B*)** Immunoblot analysis of Gnd-SPA or GndA-SPA levels. *E. coli* MG1655 strains carrying the SPA fusions were grown in LB medium under aerobic or anaerobic conditions at 30°C to mid-exponential phase and either subjected to 42°C heat shock or maintained at 30°C for 20 min. The SPA tag was detected using an α-FLAG antibody. Under some conditions, we detect a band of some higher molecular weight (GndA*-SPA) for the GndA-SPA strain. **(*C*)** In vivo degradation assay for cells expressing either Gnd-SPA and GndA-SPA. Upon reaching mid-exponential phase (OD_600∼_0.5) at 30°C, cells were exposed to heat shock at 42°C for 5 min after translation was inhibited by the addition of 30 µg/mL chloramphenicol. Samples were collected at the indicated time points and protein levels were quantified by western blot analysis. For (B) and (C), Ponceau S-stained membranes serve as loading control.

To examine GndA expression relative to Gnd, we introduced a sequential peptide affinity (SPA) tag at the end of the *gndA* or *gnd* ORFs on the chromosome in separate strains. These cells were grown at 30°C under aerobic as well as anaerobic conditions given that the 6-phosphogluconate dehydrogenase encoded by *gnd* is a central enzyme of the pentose phosphate pathway, which links respiratory metabolism and cellular redox balance through NADPH generation (18, 19). At exponential phase, half of each culture was exposed to 42°C for 20 min given the prior report of GndA heat shock induction (10). The levels of Gnd-SPA were similar across all tested conditions (Fig. 1B). In contrast, the levels of an ∼20 kDa GndA-SPA protein were elevated for cells exposed to heat shock under aerobic but not anaerobic conditions (Fig. 1B). In a more extended time course, GndA-SPA heat shock induction was observed within 5 min (Fig. S1B).

Interestingly, a second band of ∼26 kDa (labeled GndA*) could be detected under some conditions, particularly later in growth (Fig. S1B). Another AUG codon upstream in the same frame as the 36 aa GndA protein could give rise to a 54 aa protein. However, the additional 18 aa would not account for the size of the higher molecular weight band and no significant translation initiation signal is observed at this position in the ribosome profiling data. Additionally, most sequences in an alignment of the amino acid sequence of this upstream region have a stop codon (Fig. S2A).

To eliminate Gnd, GndA*, and GndA translation, we introduced mutations to selectively disrupt the start codons of each ORF. Gnd-SPA was no longer detected when the *gnd* start codon was replaced by a stop codon (*gnd*_STOP_) (Fig. S1C). To disrupt GndA without altering the Gnd amino acid sequence, we introduced three nucleotide substitutions that eliminate the three out-of-frame start codons but were silent with respect to Gnd translation (*gndA*_SILENT_) (Fig. S1C). Both GndA*-SPA and GndA-SPA levels were substantially reduced by the *gndA*_SILENT_ mutations. Interestingly however, the effects of the *gndA*_SILENT_ and *gnd*_STOP_ mutations were not reciprocal with respect to Gnd and GndA (Fig. S1C). Whereas *gndA*_SILENT_ had no detectable effect on Gnd-SPA levels, the *gnd*_STOP_ mutation substantially reduced levels of both GndA*-SPA and GndA-SPA. Possible explanations for this observation are that eliminating the *gnd* start codon affects translation in the alternate reading frame, the full-length transcript is destabilized in the absence of Gnd translation, and/or the GndA protein is less stable in the absence of the Gnd protein.

To test what sequences are required for GndA*-SPA and GndA-SPA expression, we generated a series of pBAD24 plasmids expressing GndA-SPA from the P*_araBAD_* promoter but with different sequences upstream of the 36-aa *gndA* start codon; 335 bp, 295 bp (starting with the *gnd* start codon), 250 bp, 56 bp (2 bp before the 54-aa *gndA* AUG) (indicated in Fig. 1A). We also generated a plasmid with the 36-aa *gndA* ORF fused to SPA but translated from the RBS on the plasmid. Expression of the GndA-SPA fusion alone resulted in low expression of a single band (Fig. S1D). A band corresponding to GndA-SPA could also be detected for the other constructs. Interestingly however, GndA*-SPA was only detected for the 335 bp and 295 bp constructs containing the *gnd* start codon suggesting that its production depends on this start codon (Fig. S1D). We therefore propose that the *gnd* locus supports production of two GndA-related species: GndA, whose translation is initiated from the internal *gndA* start codon and an extended form, GndA*, whose production requires the *gnd* AUG. We further suggest that translation of GndA* involves a +1 frameshift so the protein ends in the alternative *gndA* reading frame tagged with SPA. Although the precise mechanism underlying GndA* production remains to be established, our observations suggest that the upstream *gnd* sequence and internal RNA structure may together enable an unconventional mode of translation within the overlapping *gnd* locus. In this study, we focused on the protein that would initiate with the start codon giving rise to a 36 aa GndA protein, given the stronger heat shock induction.

### GndA is less stable than Gnd

To test whether Gnd impacts GndA-SPA levels, we examined GndA-SPA expressed from an arabinose-inducible P_BAD_ promoter in wild type (WT) and Δ*gnd* backgrounds (Fig. S1E). We found that GndA-SPA levels are comparable in the two backgrounds, though the levels of GndA-SPA increased substantially upon heat shock despite expression from the heterologous P_BAD_ promoter and plasmid RBS. This observation suggests that the P_BAD_ promoter is temperature-responsive and/or that GndA is sensitive to differential degradation at low and high temperatures. We also specifically assessed the stabilities of chromosomally encoded Gnd-SPA and GndA-SPA by adding chloramphenicol to inhibit translation 5 min after heat shock (Fig. 1C). We did not observe any change in Gnd-SPA levels suggesting the dehydrogenase is very stable. In contrast, there was a rapid decrease in GndA-SPA levels, indicating the smaller protein is much less stable. These results indicate that Gnd and GndA have significantly different half-lives, though the SPA tag could have some impact on proteolysis.

### Heat shock-dependent translation of GndA is controlled by a change in RNA secondary structure

We next examined how GndA expression is specifically induced by heat shock. Alternative transcription start sites and RNA processing have been proposed as mechanisms modulating internal translation in eukaryotic systems (reviewed in (20)). Thus, we first examined the *gnd/gndA* transcript in cells grown in LB medium with or without heat shock under aerobic and anaerobic conditions using a probe hybridizing to the *gndA* coding region (*gnd* codons 110–123) (Fig. S3A). For all conditions, we detected a single predominant ∼1,500 nt *gnd/gndA* transcript, expected to encode the 1,407 nt *gnd* ORF. The levels of the *gnd/gndA* transcript did not change upon heat shock under aerobic conditions, though the levels were decreased under anaerobic growth. We did not observe a *gndA*-specific transcript generated by alternate transcription or RNA processing. These results suggest that both Gnd and GndA are translated from the same mRNA.

It was previously proposed that internal base pairing within the *gnd* transcript could give rise to two alternative RNA secondary structures (14–16). One structure involves base pairing between the mRNA leader region (region I here) and sequences upstream of the *gnd* Shine–Dalgarno (SD) sequence (region IIa here). A second structure involves long-range base pairing between the SD sequence (region IIb here) and a complementary sequence located deep within the *gnd* coding region, around codons 69–78 (region III here) (Fig. 2A and S3B). This structure, termed the “internal complementary sequence (ICS)” (15), was proposed to regulate growth rate-dependent differences in 6-phosphogluconate dehydrogenase levels in minimal medium with glucose compared to acetate (21). In this latter duplex, one strand of the stem overlaps the *gnd* RBS (region IIb), while the second strand ends ∼70 nt upstream of the GndA start codon (region III). The sequences of both strands exhibit a high degree of conservation across the diverse Enterobacteriaceae genomes (Fig. 2B). We also predict that region III could base pair with another region (region IV). Each of these proposed duplexes has evidence of sequence conservation in both strands as well as predicted structural stability (Fig. 2B).

**Fig. 2.**
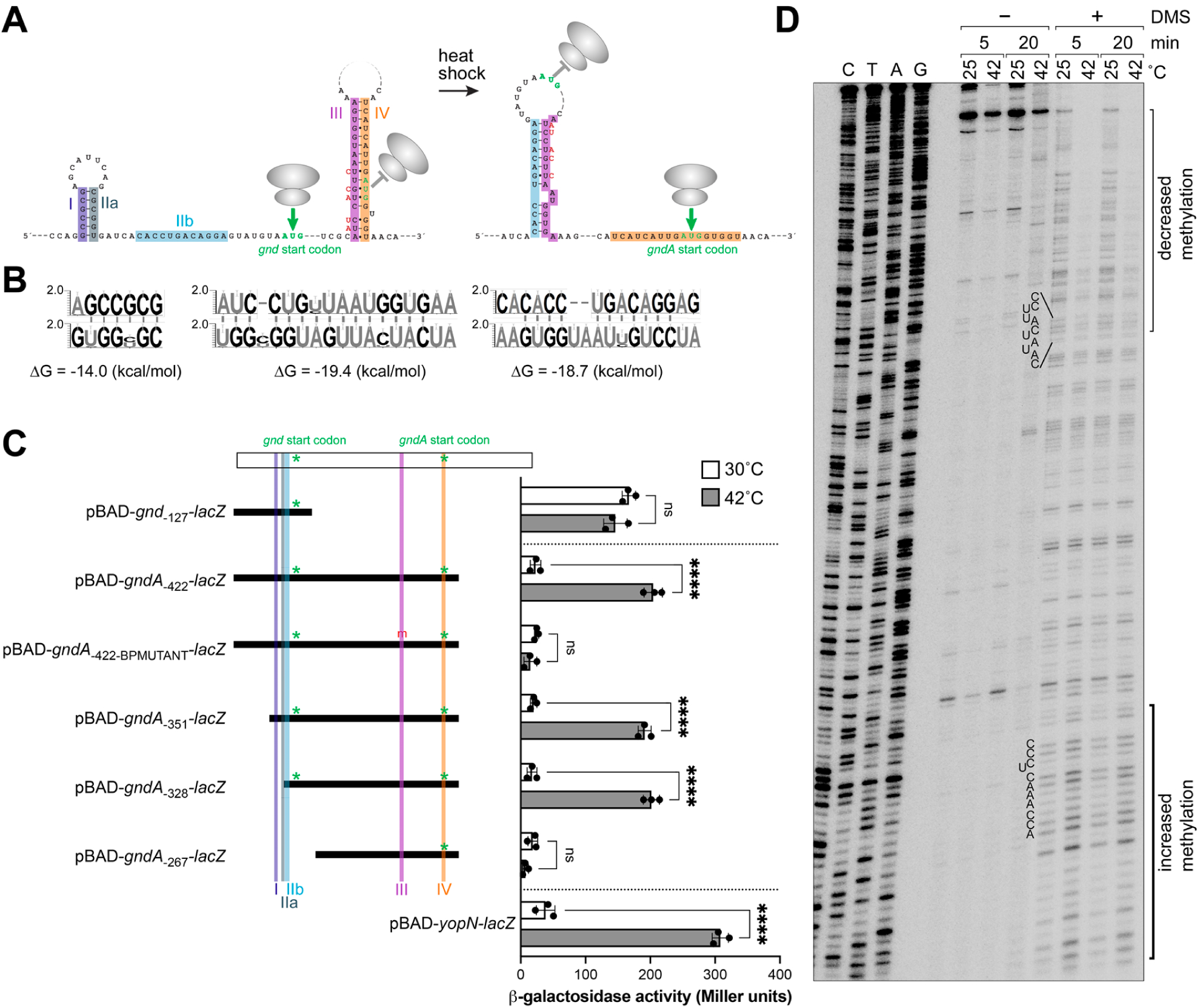
Heat shock induction involves internal complementary sequence. **(*A*)** Predicted RNA secondary structures for *gnd* transcript. Left and right structures are based on previously reported structures (16) while middle structure is predicted here. Regions I, IIa, IIb, III, and IV sequences involved in predicted base pairing are highlighted in different colors. Start codons for *gnd* and *gndA* based on ribosome profiling are in green font. Mutations to disrupt pairing with region III are in red font. **(*B*)** Sequence logos illustrate the conservation of both strands of the duplexes/stems shown in panel (A). Duplexes free energy was predicted using Afold (43). **(*C*)** β-galactosidase assays of translational fusion to *bgabg* (heat stable β-galactosidase) under control of the arabinose-inducible promotor pBAD. The fusions contain the first 10 codons of *gnd* or *gndA* fused to *bgab* and the indicated regions upstream of the respective start codons. Cells were grown in LB medium at 30°C to mid-exponential phase (OD_600∼_0.5) supplemented with 0.1% L-arabinose. Cultures were then equally divided into pre-warmed flask and incubated at 30 or 42°C for 20 min. The RNAT of *yopN* served as control (23). β-galactosidase activities are shown in Miller Units ± SD were of three independent biological replicates with statistical significance assessed by one-way ANOVA **(*D*)** In vivo DMS structure probing of *gnd* RNA at low and high temperature. Cells were grown to mid-exponential phase then equally divided into pre-warmed flask and incubated at 25 or 42°C for 20 min and treated with 50 µM DMS for 5 or 20 min at the respective temperature. RNA was isolated and used for subsequent primer extension inhibition analysis. Lanes 1-4 show the DNA sequencing ladder. Dark bracket denotes region for which modification increased, and light bracket denotes region for which modification decreased at higher temperature. Some sequence in these regions is listed. The corresponding sequences are indicated with black dots (increased modification) and white dots (decreased modification) in Fig. S3B.

The observed heat shock-dependent translation of GndA-SPA together with the potential of internal base pairing is reminiscent of the well-described translational control by temperature-responsive RNA secondary structure (RNA thermometers). At lower temperatures, stem loops typically sequester the RBS. At higher temperatures, these structures are destabilized, sometimes leading to alternative structures (reviewed in (22)). We thus hypothesized that at low temperature, region III does not form a stable duplex with region IIb, leaving region IIb largely unpaired and accessible for *gnd* translation. Instead, region III may interact with other sequences such as region IV, potentially forming a stable duplex that blocks *gndA*translation (Fig. 2A). We propose that at elevated temperatures, the RNA structure rearranges to form the ICS stem-loop between regions IIb+III, which promotes *gndA*translation by increasing access to the *gndA* RBS and/or facilitating ribosome frameshifting. The RNA structures and translation of either *gnd* or *gndA* additionally may be shaped by the interplay between transcription and translation.

To test if heat shock induced translation of GndA is dependent on the sequences encompassing the ICS secondary structure, we generated various translational reporter fusions between the first 10 codons of either *gnd* or *gndA* and *lacZ* encoding a temperature-resistant enzyme (*bgabg*) and measured β-galactosidase activity at 30 and 42°C (Fig. 2C). The *gnd-lacZ* fusion with 127 nt upstream of the *gnd* start codon did not show heat shock induction, while we observed strong heat-shock induction of *yopN-lacZ*, a control fusion to a characterized RNA thermometer (23). For the *gndA-lacZ* fusions with 422-nt (same 5’ end as *gnd-lacZ* fusion) or 351-nt (starting with prominent transcription start side), which both contained regions I-IV and the first 10 codons of *gndA*, we observed clear heat-shock-dependent induction. A construct starting 328-nt upstream of the *gndA* start, which lacked the region I-IIa sequences, also was clearly induced. However, a fusion with only 267-nt, which lacked region IIb, eliminated the heat shock induction of the *gndA-lacZ* fusion. Similarly, in the context of the 422-nt fusion, mutations in region III that are silent with respect to the *gnd* CDS but were predicted to disrupt base pairing between regions II-III (*gndA*_BPMUT_) also eliminated the heat shock induction. These results suggest that ICS pairing between regions IIb-III is required for temperature-responsive induction of GndA translation.

To assess RNA secondary structure changes within the *gnd* transcript in vivo, cells grown at 25°C or at 42°C were treated with 50 mM dimethyl sulfate (DMS) for 5 min, while control samples were treated with water. DMS preferentially methylates the N1 position of adenine and the N3 position of cytosine in single-stranded RNA. Total RNA was isolated, and DMS-induced modifications were detected by primer extension analysis of the *gnd* transcript (Fig. 2D). A comparison of the 25°C and 42°C samples treated with DMS revealed that segments of the *gnd* RBS and the ICS are partially protected (thin bracket) from chemical modification in vivo under heat shock conditions suggesting increased secondary structure. In contrast, positions immediately upstream of *gndA* (thick bracket) display increased DMS reactivity upon heat shock consistent with decreased secondary structure. Together, these results demonstrate that the *gnd* mRNA adopts distinct structural states at low and high temperature, likely facilitating translation of the *gndA* altORF.

### GndA is needed for heat shock survival

While the *E. coli* heat shock response has been well characterized (reviewed in (24)), the contribution of the *gnd* locus to heat shock survival has not been examined. Thus, we asked whether deletion of the complete locus or functional disruption of *gnd* and *gndA* (Fig. 3A) affected survival under heat shock conditions. The corresponding strains were grown in LB medium at 30°C to mid-exponential phase, exposed to heat shock at 42°C for 20 min, whereupon survival was quantified by determining colony-forming units (CFU) (Fig. 3B). Deletion of the complete *gnd* coding sequence (*Δgnd*) resulted in a pronounced reduction in CFU recovery compared to the WT strain. This reduction was not observed for a strain expressing Gnd-SPA, suggesting tagged Gnd is functional. In contrast, cells with the GndA-SPA fusion, in which the 3’ end of the primary *gnd* gene is disrupted, were sensitive to heat shock. Similarly, disruption of *gndA* translation with the *gndA*_SILENT_ mutations that leave Gnd-SPA expression intact, resulted in a significant reduction in CFU recovery following heat shock. Thus, both the primary and alt-ORF gene products are needed to protect against higher temperature (Fig. 3B). Consistent with our results, a preprint reported reduced CFU formation following heat shock (45°C) for *gnd* and *gndA* mutants (25).

**Fig. 3.**
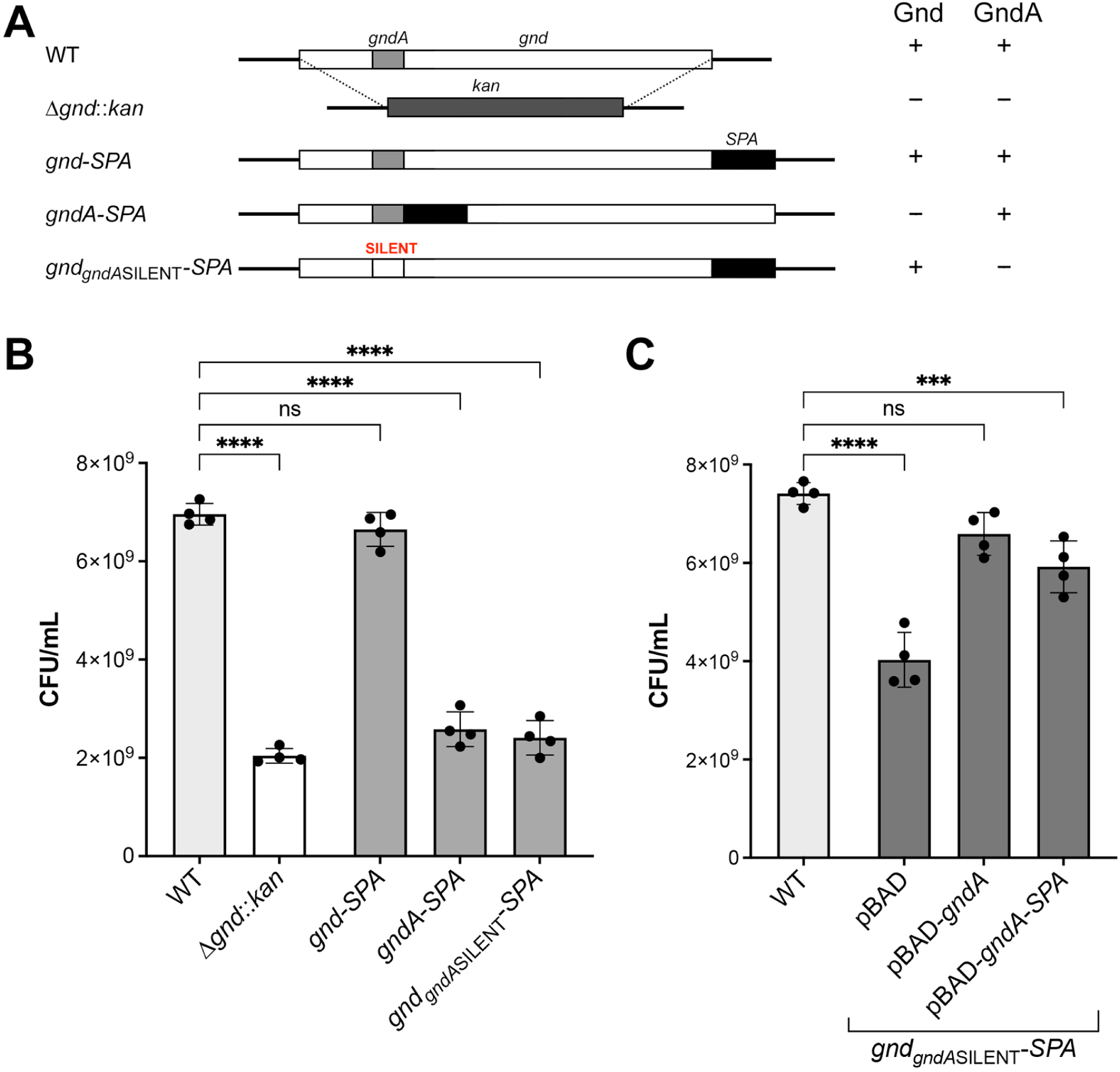
GndA is critical for heat shock resistance. **(*A*)** Diagram of mutants and expected expression of Gnd and GndA. **(*B*)** Colony-forming unit (CFU) formation following heat shock for cells lacking Gnd, GndA or both. **(*C*)** Trans-complementation of heat shock resistance in *gnd*-*_gndA_*_SILENT_-*SPA* strain lacking GndA. The *gnd*-*_gndA_*_SILENT_-*SPA* strain mutations harbored either the empty vector (pBAD) or pBAD expressing GndA or GndA-SPA. Expression from pBAD was induced 15 min prior to heat shock by addition of 0.2% arabinose. For (B) and (C), cells were grown at 30°C to mid-exponential phase (OD_600∼_0.5) and transferred to pre-warmed flasks for heat shock exposure (42°C, 20 min). CFUs were determined following overnight recovery. Error bars represent the mean ± SD for four independent biological replicates with statistical significance assessed by one-way ANOVA.

We also tested whether untagged and tagged GndA expressed from a P_BAD_ plasmid could complement the *gndA*_SILENT_ mutant strain (Fig. 3C) and found that pBAD-GndA restored almost WT growth while pBAD-GndA-SPA partially complemented the growth defect. Together, these findings support a model in which heat shock survival depends on GndA.

### GndA-SPA co-fractionates with the membrane

To gain insights into the role of the GndA protein, we carried out structural predictions. AlphaFold3 (26) suggests GndA adopts a predominantly α-helical conformation and helical wheel analysis (27) suggests an amphipathic organization, with hydrophobic residues on the opposite face of the helix relative to polar and charged residues (Fig. S4). In line with these predictions, GndA was initially identified in a membrane proteomics screen (10), and GndA overexpressed from a plasmid was reported to associate with the inner membrane (25). Following subcellular fractionation of cells grown in LB medium, we also detected chromosomally encoded GndA-SPA in the membrane fractions (like the outer membrane control protein OmpC) (Fig. 4A).

**Fig. 4.**
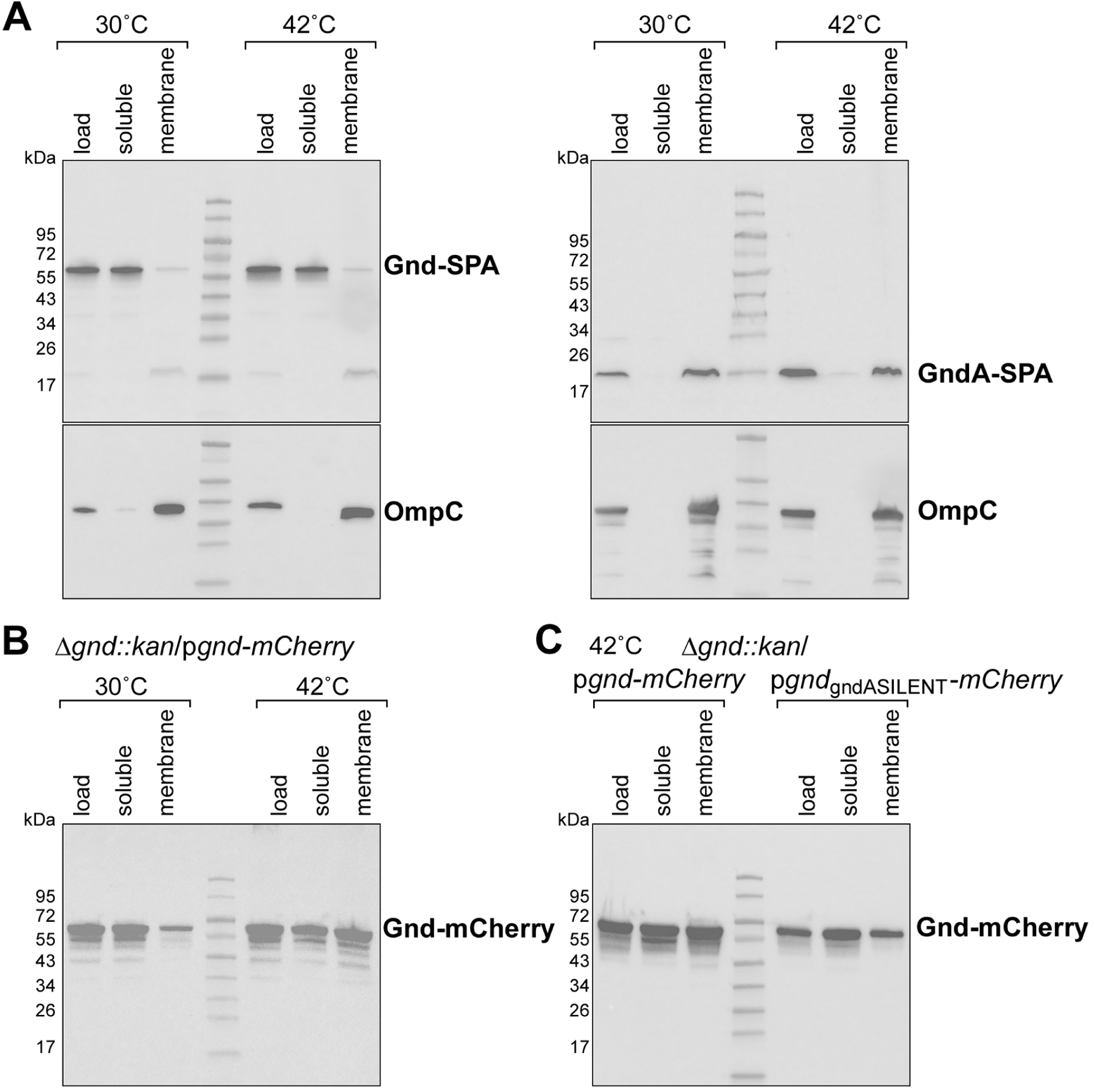
GndA-SPA co-fractionates with the membrane. **(*A*)** Subcellular fractionation of proteins for the Gnd-SPA (left) and GndA-SPA (right) strains. The upper panel shows SPA-tagged proteins detected using an α-FLAG antibody. As a fractionation control, the lower panel shows the outer membrane protein OmpC detected with an α-OmpC antibody. **(*B*)** Subcellular fractionation of proteins for Gnd-mCherry for cells grown without and with heat shock. **(*C*)** Subcellular fractionation of proteins for Gnd-mCherry without (left) and with (right) the gndA_SILENT_ mutation strains for cells exposed to heat shock. For both (B) and (C), the respective strains were grown at 30°C in LB medium to mid-exponential phase (OD_600_∼0.5). The culture was split, with one half subjected to heat shock at 42°C for 20 min and the other maintained at 30°C. For (D), both cultures grown at 30°C in LB medium to mid-exponential phase (OD_600_∼0.5) were subjected to 42°C for 20 min. Cells were fractionated into cytosolic (C, soluble) and membrane (M) fractions and compared to the whole-cell lysate (L).

In contrast to GndA-SPA, the larger Gnd-SPA protein was predominantly observed in cytosolic fractions for cells grown at both 30 and 42°C, although intriguingly, a small fraction of Gnd-SPA was detected in the membrane fraction (Fig. 4A). Given the observation that Gnd-SPA is very stable, we also examined fractionation of Gnd C-terminally tagged with mCherry after 15 min induction from an IPTG-responsive promoter on a low-copy plasmid in a Δ*gnd* background. This fractionation experiment for cells grown at low and high temperature showed that while Gnd-mCherry was detected in the cytosolic fraction for the cells grown at both temperatures, a significantly higher proportion was in the membrane fraction for the cells grown that 42°C (Fig. 4B). This redistribution of Gnd-mCherry was decreased when we introduced silent mutations in the *gnd* coding sequence that selectively abolish GndA translation (Fig. 4C). These observations indicate that the altORF-encoded GndA protein is membrane-associated and influences localization of the primary ORF-encoded Gnd protein at higher temperatures.

### Gnd is recruited to cell poles in a GndA-dependent manner upon heat shock

To visualize the subcellular localization of Gnd in individual cells, we performed fluorescence microscopy for Δ*gnd* cells expressing Gnd-mCherry from the low-copy plasmid used in the fractionation assays. Cells grown to OD_600_ ∼0.4 at 30°C were treated with IPTG for 15 min and then washed to remove the inducer. Imaging of these cells showed that Gnd-mCherry had a diffuse cytosolic localization. However, following a shift to 42°C, there was a pronounced spatial relocalization of Gnd-mCherry to foci in the polar regions of the cell (Fig. 5A). This shift was observed as early as 5 min after temperature upshift. Quantification of this response showed that 87.2% of cells displayed polar Gnd-mCherry foci after 5 min at 42°C, increasing to 92.1% after 10 min and 95.7% after 15 min. Expression of mCherry alone from a high-copy plasmid did not result in membrane or polar enrichment, ruling out tag-dependent effects (Fig. S5A). To again assess whether this relocalization depends on the altORF-encoded GndA, we also examined the effects of the *gndA*_SILENT_ mutations in the *gnd* coding sequence. For this construct, heat shock-induced membrane and polar relocalization of Gnd-mCherry was no longer observed (Fig. 5A).

**Fig. 5.**
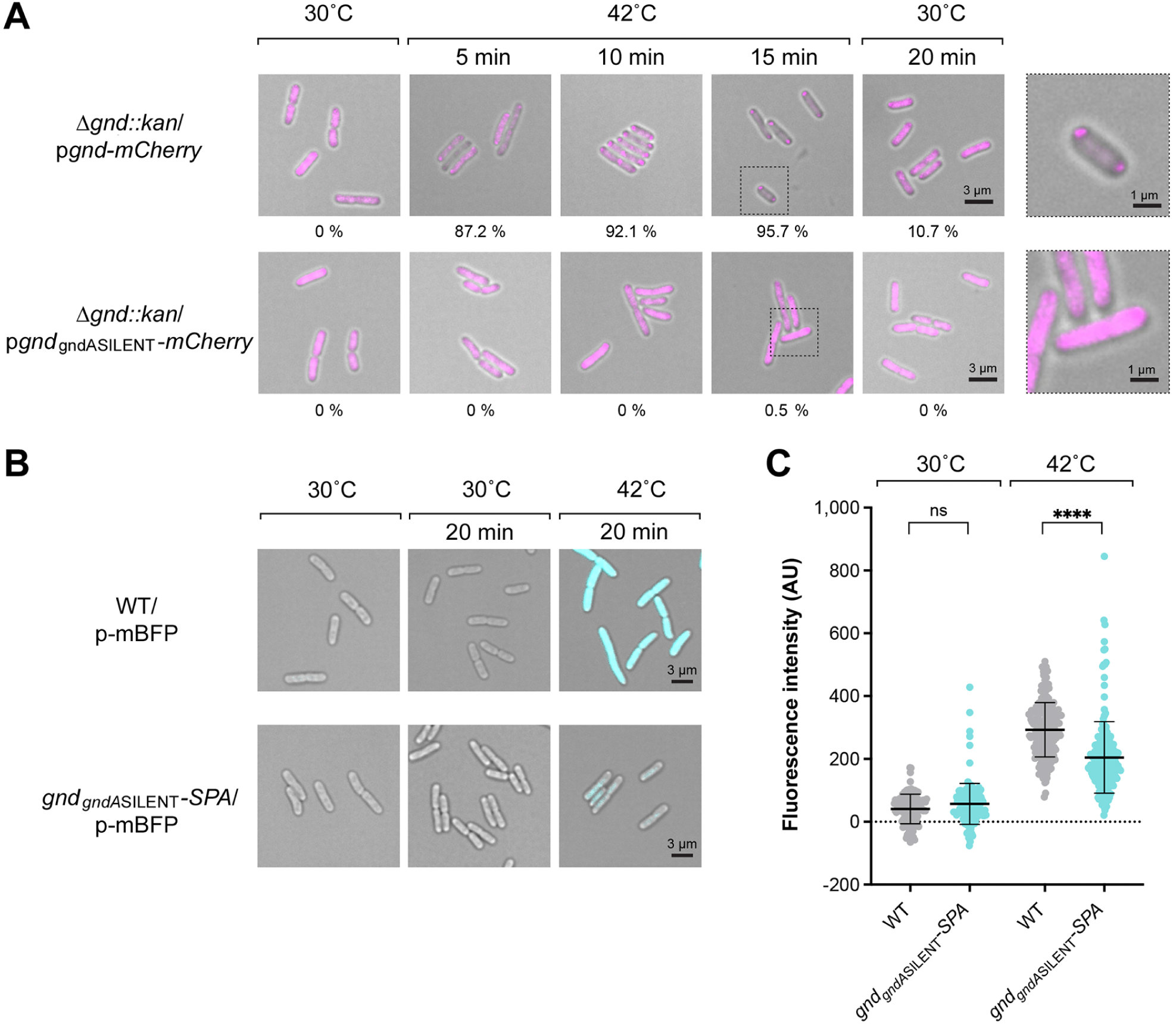
Gnd is recruited to cell poles in a GndA-dependent manner upon heat shock. **(*A*)** Representative fluorescence microscopy images of *E. coli* BL21 Δ*gnd* cells carrying a plasmid-borne *gnd-mCherry* fusion or *gnd*_gndASILENT_*-mCherry* fusion showing increasing polar localization with time after heat shock and decreasing polar localization with time after return to 30°C. Cells were grown in LB medium supplemented with glucose at 30°C to early exponential phase (OD_600_∼ 0.35). Expression was induced with 0.5 mM IPTG for 15 min. Cells were then collected and washed to remove the inducer. One fraction was resuspended in 30°C LB and a sample was imaged. The other fraction as resuspended in 42°C LB, and samples were taken at 5, 10 and 15 min after this heat shock. Subsequently, the culture was shifted back to 30°C and samples were taken at 20 min. Scale bar for left four columns = 3 µm. Polar localization for these samples was quantitated for >300 cells for each sample and the percent cells with polar localization is indicated below each image. Right column shows enlargement of cells in regions boxed by dotted line. Here scale bar = 1 µm. (***B***) Representative fluorescence images of *E. coli* MG1655 constitutively expressing mBFP. Cells were grown in LB medium supplemented with 1% glucose at 30°C to mid-exponential phase, and cultures were divided into two aliquots. One aliquot was maintained at 30°C; the other was transferred to prewarmed flasks and exposed to heat shock at 42°C for 20 min. (***C***) Quantification of mBFP fluorescence in *E. coli* MG1655 WT and *gnd_gndA_*_SILENT_-*SPA* cells under the conditions described in (B). Error bars indicate ± mean SD of n≥100 cells/strain. Statistical significance across groups was assessed by one-way ANOVA (p < 0.0001).

Since expression of the Gnd-mCherry fusion was stopped by inducer washout before the imaging and the tagged protein again showed diffuse cytosolic localization after cells were returned to 30°C for 20 min (Fig. 5A), we do not think the heat-shock induced polar localization is due to protein unfolding and aggregation. However, to more directly test whether polar Gnd-mCherry indeed relocalizes, we followed the localization of the protein at shorter time points after the shift from 42°C to 30°C. Following 15 min at 42°C, 89.9% of cells retained polar Gnd-mCherry foci 5 min after the shift back to 30°C. This fraction decreased to 55.2% after 10 min and to only 10.7% after 20 min at 30°C (Fig. S5B). In contrast, cells expressing the *gndA*_SILENT_ variant showed diffuse Gnd-mCherry localization throughout the time course, consistent with the inability of Gnd to form heat shock-induced foci in the absence of GndA. Together, these results show that GndA is required for heat shock-responsive spatial relocalization of Gnd to the membrane poles, which is reversed following recovery from heat shock.

The Gnd dehydrogenase is a central hub for cellular NADPH production in *E. coli* (28). Given the GndA-dependent spatial relocalization of Gnd during heat shock, we asked whether heat shock and mutations in GndA affect intracellular NADPH dynamics. We used the metagenomic-derived blue fluorescent protein (mBFPt) sensor (29), expressed from a constitutive promoter to monitor free NADPH levels. NADPH binding by mBFP rapidly enhances the intrinsic blue fluorescence of the NADPH cofactor, enabling real-time monitoring changes in intracellular levels (30). We first examined mBFP fluorescence in WT *E. coli* MG1655 cells. The WT cells grown at 30°C exhibited little to no detectable blue fluorescence, in contrast to cells exposed to 42°C for 20 min (Fig. 5B). To assess whether GndA contributes to the increased fluorescence at 42°C, we compared WT cells with cells carrying the chromosomal *gnd_gndA_*_SILENT_ mutations that silence GndA expression (Fig. 5B). Quantification of n≥100 cells per condition showed both strains had similarly low fluorescence at 30°C. However, following heat shock, fluorescence increased 7.1-fold in the WT, while fluorescence in the *gnd_gndA_*_SILENT_ cells only increased 3.6-fold, significantly lower than for the WT cells (Fig. 5C). These results suggest a potential link between GndA-dependent relocalization of Gnd and free NADPH levels.

## Discussion

Bacterial adaptation to rapidly changing environmental conditions requires regulatory mechanisms that enable fast and efficient proteome remodeling. AltORF-encoded proteins, translated from positions internal to annotated coding sequences, represent a largely unexplored layer of bacterial coding capacity that may expand the functional versatility of the bacterial proteome. Here, we report that an altORF-encoded microprotein is selectively translated during heat shock from the same transcript as its primary ORF partner, *gnd*, encoding 6-phosphogluconate dehydrogenase. We show this selective translation is mediated by a thermoresponsive RNA secondary structure that promotes altORF expression upon heat shock. Furthermore, we uncover a functional role for the altORF-encoded GndA microprotein in recruiting the primary ORF-encoded dehydrogenase to the cell poles during heat stress. Together, our results reveal a previously unrecognized mechanism by which bacteria coordinate a stress-response from two proteins encoded in different frames by the same sequence.

### Translational regulation by ORF-internal RNA secondary structures

RNA structures, including intrinsic RNA secondary structures and regulatory RNA-mRNA base pairing, are important determinants of post-transcriptional regulation in bacteria (reviewed in (31, 32)). To date, the focus for riboregulation has been on the 5’ untranslated regions (5’-UTRs) of annotated coding sequences. For example, temperature-induced RNA structural remodeling in 5’ UTRs has been shown to modulate translation of multiple mRNAs (reviewed in (22, 33)). Here we show that RNA structural elements within a coding sequence can also modulate translation. We suggest that translation of the altORF GndA is conditionally induced during heat shock through a temperature-sensitive remodeling of secondary structure of the *gnd* transcript. It was similarly found that RNA folding within the *petH* transcript of the cyanobacterium *Synechocystis* controls internal translation, generating two protein isoforms with opposing activities (34). Condition-dependent RNA structural remodeling internal to coding sequences may be far more prevalent than previously appreciated and serve as a signature for identifying translated altORFs.

### Membrane localization of 6-phosphogluconate dehydrogenases

Using fusion constructs, we observed GndA-mediated Gnd redistribution to the membrane following heat shock, with relocalization detectable as early as 5 min after stress onset. Although 6-phosphogluconate dehydrogenase is typically cytoplasmic, proteomic analyses of *Streptococcus pneumoniae* detected the enzyme in cell wall-associated fractions (35). Additionally, a second family of short-chain 6-phosphogluconate dehydrogenases has been described (36). For the *Gluconacetobacter diazotrophicus* enzyme, which converts NAD⁺ to NADH, the active homodimeric enzyme is detected in both soluble and membrane fractions. It was reported that this membrane-associated 6-phosphogluconate dehydrogenase supplies NADH directly to membrane-bound NADH dehydrogenases, thereby coupling carbohydrate oxidation to respiratory energy conservation. By analogy, the membrane localization of *E. coli* Gnd in response to heat shock could position NADPH production close to sites of high redox demand at the cytoplasmic membrane where reactive oxygen species are generated by respiratory activity (18). In addition, or alternatively, relocalization of pentose phosphate pathway enzymes to the membrane could facilitate efficient capture and processing of imported glucose derivatives, particularly under conditions requiring rapid metabolic reprogramming. Thus, membrane re-localization of Gnd after heat shock could position the enzyme near sites of greatest NAD need as well as substrate entry. Such an arrangement could promote efficient NADPH generation while minimizing diffusion between transport, metabolism, and membrane-associated redox processes.

### Protein recruitment to membranes by small amphipathic helix proteins

Although bacterial cells lack membrane-bound organelles, proteins are not randomly distributed but are often targeted to distinct regions even within the cell membrane, where different regions can have different curvatures (reviewed in (37)). Two other amphipathic helix proteins in addition to the GndA protein have been implicated in recruiting proteins to membranes. The first characterized example is the 26-aa *Bacillus subtilis* SpoVM protein, which localizes to the forespore membrane during endospore formation (38). A second example is the 28-aa *E. coli* AzuC protein, which is localized to the membrane throughout the cell, and like GndA promotes membrane association of a dehydrogenase, in this case, the aerobic glycerol-3-phosphate dehydrogenase (39). AzuC co-purifies with the dehydrogenase, promoting its membrane association and enhances its activity. Intriguingly, SpoVM, AzuC and GndA all have proline residues at the N- or C-terminal ends. A proline in SpoVM has been shown to be critical for selective binding to positively-curved forespore membrane (38, 40) raising the question of what GndA features target the protein to the cell pole membranes, which have negative curvature. In general, we suggest that small, α-helical proteins may serve a wide-spread role as membrane recruitment entities.

### Evolution of ORF internal genes

The *gndA* coding sequence overlaps codons 99-136 of Gnd. This region lies within the linker connecting the two functional domains of the Gnd protein, a segment that is would be expected to be tolerant to sequence variation but also includes N102, an NADP-interacting residue that undergoes repositioning during the coenzyme-induced conformational change of Gnd and residues 128–130, which contribute to binding of the 6-phosphogluconate substrate (41). The overlapping arrangement therefore requires the same nucleotide sequence to accommodate the coding requirements of two proteins in different reading frames. The conservation of *gndA* within the larger *gnd* gene is consistent with selective constraints acting on the embedded protein-coding sequence (Table S1), in contrast to the ORF-internal *ftsO* small RNA encoded within *ftsI* (42), whose conservation reflects selection on RNA sequence and/or structure rather than on an alternative protein product. Deeper analyses of evolutionary constraints within ORF-internal sequences, particularly when combined with transcriptomic and ribosome-profiling data, may reveal additional functional gene products embedded within annotated protein-coding genes.

## Materials and Methods

### Strains, Plasmids and Oligonucleotides

Bacterial strains, plasmids, and oligonucleotides used in this study are listed in Dataset S1. Detailed descriptions of strain construction, plasmid generation are provided in the SI Appendix.

### Bacterial Growth

Bacterial growth was carried out in Lysogeny Broth (LB) medium at indicated temperatures and supplemented with appropriate antibiotics, described in detail in the SI Appendix.

### Immunoblot and Northern Analysis

Specific tagged proteins were detected by immunoblotting, and target RNAs were analyzed by northern blotting, as described in detail in the SI Appendix. For select experiments, proteins were isolated from subcellular fractions prior to analysis.

### Microscopy

Bright-field and fluorescence microscopy of bacterial cells was carried out according to protocols detailed in the SI Appendix.

### β-Galactosidase Activity Assays

Activity of *lacZ* reporter constructs corresponding to *gnd* and *gndA* was measured as described in detail in the SI Appendix.

### In vivo DMS structure probing

In vivo structure probing of RNA secondary structure under varying temperature conditions was performed as described in detail in the SI Appendix.

### Bioinformatic Analysis

RNA folding prediction, analyses of species distribution, phylogeny, and genomic context were conducted as detailed in the SI Appendix.

## ACKNOWLEDGMENTS.

We thank A. Buskirk for generating browser images, F. Narberhaus (Ruhr University Bochum) for the pBAD-*bgaB*-His and pBAD2-yopN-bgaB-His plasmids and K. Ramamurthi for supporting the microscopy carried out in his lab. We appreciate the helpful experimental and manuscript suggestions of F. Naberhaus, K. Ramamurthi and members of the Storz and P. Adams labs. Work by J.J.S. was supported by an NICHD Early Career Development Award. This research was supported by the Intramural Programs of the *Eunice Kennedy Shriver* National Institute of Child Health and Human Development (J.J.S., A.Z., R.Z. and G.S.), the of the Center for Cancer Research of the National Cancer Institute (F.R.-L.) and the National Library of Medicine (S.A.S.).

