## Supporting Information for "AltORF-encoded microprotein recruits primary-ORF-encoded dehydrogenase to cell poles upon heat shock"

**SI Appendix**

**Schmidt et al.**

**Strains, Plasmids and Oligonucleotides.** A detailed list of all plasmids, oligonucleotides and bacterial strains used in this study are summarized in **Dataset S1**. Oligonucleotides were purchased from Integrated DNA Technologies and gene fragments were synthesized by Twist Bioscience, respectively. The *gnd* (b2029) and *gndA* genes were tagged on the chromosome, as previously described (1) using the λ red recombination system in *E. coli* NM400 or *E. coli* NM500 by replacement of the stop codon of each gene with the mutation linked to a SPA-kanamycin or SPA-chloramphenicol cassette. Constructs were then moved into *E. coli* MG1655 by phage transduction using a lab stock of P1 phage. Similarly, gene deletions of *gnd* and *lacZ* (b0344) were generated by replacing each coding sequence (CDS) with a kanamycin cassette, followed by P1-mediated transduction into *E. coli* MG1655. Cloning was performed using Q5® High-Fidelity 2X Master Mix (New England Biolabs, #M0492L), NEBuilder® HiFi DNA Assembly Master Mix (New England Biolabs, E2621S) and NEB® 5-alpha Competent *E. coli* (New England Biolabs, #C2984H). Start to stop mutants of *gnd* and *gndA* were synthesized by Twist Bioscience and introduced chromosomally as previously mentioned. Fusions to *bgaB* were constructed using standard restriction and ligation protocols using NheI and EcoRI restriction sites on the pBAD-*bgaB-his* vector backbone (2). Restriction enzymes and buffers were purchased from New England Biolabs (#R3131S; #R3101S; #B6004S). GndA-SPA fusions expressed from an arabinose-induced promoter were constructed using NEBuilder® HiFi DNA Assembly Master Mix or standard restriction with SmaI (New England Biolabs #R0141S) and ligation into pBAD24. The sequences of all constructs were verified by nanopore sequencing (Plasmidsaurus) or colony PCR using *Taq* 2X Master Mix (New England Biolabs, #M0270) followed by Sanger sequencing (Azenta Life Sciences).

**Bacterial Growth.** Bacterial strains were cultivated in Lysogeny Broth (0.5% (w/v) yeast extract, 1% (w/v) tryptone, and 1% (w/v) NaCl; LB) medium supplemented with the appropriate antibiotics: kanamycin (50 µg/mL), chloramphenicol (50 µg/mL), ampicillin (100 µg/mL), or spectinomycin (30 µg/mL). Unless otherwise indicated, bacterial cultures were grown at 30˚C with shaking at 250 rpm. Here, heat shock growth conditions are defined as incubation at 42˚C.

**Immunoblot Analysis.** For protein extraction, cells from 1.0 mL of cell culture were collected and resuspended in 1X phosphate-buffered saline (PBS) at a concentration corresponding to an $OD_{600}$ of 1 per 80 µL. Cell suspensions were mixed with 2x Laemmli sample buffer (Bio-Rad, #1610737), heated at 95 °C for 10 min, and centrifuged at 4 °C for 3 min. The corresponding supernatant was analyzed by SDS–PAGE (Bio-Rad, #4561093; 4–20%) and transferred onto nitrocellulose membranes (Invitrogen). Total protein content was visualized using Ponceau S staining (Sigma-Aldrich #6226-79-5) and imaged with the ChemiDoc™ MP Imaging System (Bio-Rad). Ponceau S-stained membranes were subsequently washed with 1X PBST (PBS containing 0.1% Tween-20). Membranes were probed with anti-FLAG (M2)-HRP conjugate (Sigma-Aldrich, #A8592-1MG) or anti-mCherry-HRP conjugate (Invitrogen #740003TH2P) antibodies following the manufacturer's instructions and imaged using the ChemiDoc™ MP Imaging System.

**Northern Blot Analysis.** Total RNA was isolated using TRizol (Thermo Fischer Scientific) as described previously (3). RNA was resuspended in 20–50 µL of DEPC-treated $H_2O$ and quantified using a NanoDrop spectrophotometer (Thermo Fisher Scientific). Total RNA (5-10 µg per lane) was separated on a 8% denaturing polyacrylamide gels (shorter RNAs) or agarose gel (longer RNAs). The polyacrylamide gels were prepared with 6 M urea (1:4 mix of Ureagel Complete to Ureagel-8 from National Diagnostics) with 0.08% ammonium persulfate and run in 1X TBE buffer at 300 V for 90 min. RNA was transferred to a to a Zeta-Probe GT membrane (Bio-Rad) at 20 V for 16 h in 0.5X TBE. Agarose gels were prepared with 2% NuSieve 3:1 agarose (Lonza), 1X MOPS and 2% formaldehyde and RNA was transferred to a Zeta-Probe GT membrane (Bio-Rad) via capillary action overnight. For both blot types, RNA was immobilized on the membranes by UV crosslinking. RiboRuler High- and Low-Range RNA ladders (Thermo Fisher Scientific) were visualized by UV shadowing. Membranes were then blocked in ULTRAhyb-Oligo Hybridization Buffer (Ambion) and hybridized with 5′-$^{32}$P-end-labeled oligonucleotide probes (listed in **Dataset S1**). Following overnight hybridization, membranes were washed sequentially with SSC/0.1% SDS (2 times) and 0.2x SSC/0.1% SDS (1 time) before being exposed to X-ray film. To re-use membranes, blots were stripped by 7 min washes in boiling 0.2% SDS for 7 min (2 times), followed by 7 min washes in boiling water (2 times).

**β-Galactosidase Activity Assays.** *E. coli ΔlacZ* cells were transformed with plasmids carrying either gnd- or *gndA-bgaB-his* fusions. Cultures were grown in LB medium supplemented with ampicillin at 30°C to an $OD_{600}$ of 0.5, followed by induction with 0.1% (final concentration) L-arabinose. The cultures were then divided, with one half shifted to 42°C and the other maintained at 30°C for an additional 30 min. Samples were collected for β-galactosidase measurements, and $OD_{600}$ values were recorded simultaneously. For each assay, reaction buffer was freshly prepared by combining 10 mL of Z buffer (60 mM $Na_2HPO_4$, 40 mM $NaH_2PO_4$, 10 mM KCl, 1 mM $MgSO_4$) with 2.7 mL of β-mercaptoethanol. Reaction buffer (400 µL0 and chloroform (30 µL) were added to 400 µL of culture in a 1.5 mL tube. Samples were vortexed for 30 sec and incubated at 55˚C for 15 min. The reaction was initiated by adding 400 µL of o-nitrophenyl β-D-galactopyranoside (ONPG; 2 mg/mL, Sigma-Aldrich #N1127-5G) and incubated at 55˚C for 5–60 min. Reactions were stopped by adding 400 µL of 1 M $Na_2CO_3$, followed by centrifugation at 14,000 rpm for 5 min. The $A_{420}$ of the supernatant was measured and β-galactosidase activity was calculated in Miller units as described previously (4). Results are reported as mean ± standard deviation (SD) from three independent biological replicates. *bgaB-his* fused to the short 5′-UTR of *yopN* (*Yersinia pseudotuberculosis*) and 30 nt of its coding sequence was used as positive control for RNA-thermometer activity (5).

**In vivo Structure Probing.** Cells were grown in LB medium at 25˚C to mid-exponential phase ($OD_{600}$≈ 0.5) in a total culture volume of 250 mL. Cultures were divided into four 20 mL aliquots in Falcon tubes, with two samples subjected to heat shock at 42˚C and two maintained at 25˚C. One sample for each condition was treated with 50 mM DMS, while the other received an equal volume of water as a control. DMS treatment was carried out for 5 min and reactions were immediately quenched by addition of 0.7 M β-mercaptoethanol, followed by centrifugation at 4˚C for 20 min. Supernatants were removed and cell pellets were flash-frozen in liquid $N_2$. Total RNA was isolated using TRIzol (Thermo Fisher Scientific) as previously described (3). RNA samples were subjected to primer extension analysis using the 5′-$^{32}$P-end-labeled oligonucleotide JSP94 and AMV reverse transcriptase (AMV007-1; Life Sciences) together with the supplied reaction buffer (ARB45). Reactions were terminated by addition of 6X formamide loading dye. The same primer and was used for the corresponding sequencing reactions, using a 250 pmol PCR template generated via oligonucleotides JSP50 + JSP51 (listed in **Dataset S1**). Sequencing ladders were generated using the SequiTherm EXCEL™ II DNA Sequencing Kit according to the manufacturer's instructions. Reaction products were resolved on an 8% polyacrylamide gel and visualized by autoradiography.

**Heat Shock Survival Assay.** WT *E. coli* cells or the corresponding mutant variants were grown in LB medium to an $OD_{600}$ of 0.4-0.6. Cultures were split, with one half exposed to heat shock at 42˚C for 20 min and the other maintained at 30˚C. Cells were normalized to an $OD_{600}$ of 0.1 in fresh LB medium and survival was assessed by plating 100 µL of appropriate dilutions per plate and counting colony-forming units (CFUs). Data are presented as ± mean standard deviation (SD) from three independent biological replicates, each with three technical replicates.

**Sub-cellular Fractionation.** Cells carrying chromosomally encoded Gnd-SPA, GndA-SPA, or the pHM-plasmids encoding Gnd-mCherry or $Gnd_{GndASILENT}$-mCherry were grown in LB medium to an $OD_{600}$ of 0.4–0.6. Cultures were then divided, with one half subjected to heat shock at 42°C for 20 min and the other maintained at 30˚C. Cells were immediately harvested by centrifugation at 10,000 x g for 10 min at 4˚C. Cell pellets were resuspended in fractionation buffer (10 % w/v glycerol, 150 mM NaCl, 50 mM Tris-HCl pH 7.5) and one tablet cOmplete™ Protease Inhibitor Cocktail (Roche). The cell suspension was lysed via three passes at 20,000 psi through a microfluidizer processor (Mircrofluidics), followed by centrifugation of the crude lysate at 15,000x g for 30 min at 4˚C. Cleared lysates where then centrifuged at 130,000 g for 2.5 h at 4ºC in a TLA100.3 rotor (Beckman Optima TLX tabletop centrifuge) for separation of the soluble and membrane fraction. Following centrifugation, the top layer (soluble fraction/cytosolic fraction) was removed. The pellet was washed 3 times in fractionation buffer and finally resuspended in 500 µL fractionation buffer (membrane fraction). 6X Laemmli sample buffer (Bio-Rad, #1610737) was added to all fractions (lysate, soluble fraction, membrane fraction) and assayed by immunoblotting.

**Fluorescence Microscopy.** To visualize mCherry fluorescence, strains of interest were grown overnight at 30˚C in LB broth supplemented with 30 µg/mL spectinomycin. The following morning, overnight cultures were diluted 1:100 into fresh LB broth containing 30 µg/mL spectinomycin and grown at 30˚C with shaking in a water bath to an $OD_{600}$ of approximately 0.4. Expression of the Gnd-mCherry fusion protein was then induced by addition of IPTG to a final concentration of 1 mM, and cultures were incubated for an additional 15 min at 30˚C. Following induction, cells were harvested by centrifugation at 1,500 g for 5 min and resuspended in LB broth pre-warmed to 30˚C. An aliquot was immediately collected for imaging and used as the pre-heat-shock (0 min) sample. The remaining culture was transferred to a flask pre-warmed to 42˚C and incubated at 42˚C with shaking. Samples were collected after 5, 10, and 15 min of incubation at 42˚C. Following the 15-min incubation at 42˚C, the culture was rapidly transferred to a water bath set to 30˚C and incubated with shaking to monitor recovery from heat shock. Samples were collected after 5, 10, and 20 min of recovery at 30˚C. For fluorescence microscopy, cells from each time point were mounted on 1% (w/v) agarose pads prepared in PBS and imaged using a DeltaVision Core microscope system (Applied Precision/GE Healthcare) equipped with a Photometrics CoolSnap HQ2 camera. The data were deconvolved using SoftWorx software as described previously (6). Images were analyzed using Fiji.

To visualize the NADPH-responsive fluorescent reporter mBFP, strains harboring pCN-mBFP (7) were grown in LB medium supplemented with 1% (w/v) glucose at 30˚C to mid-exponential phase. Cultures were then divided into two equal volumes; one was maintained at 30˚C and the other was transferred to a flask prewared at 42˚C to induce heat shock. After 20 min, samples from both conditions were collected and imaged using a DeltaVision Core microscope system (Applied Precision/GE Healthcare) as indicated above. Acquisition settings were identical for all samples to allow direct comparison of fluorescence intensities. Individual cells were segmented using Cellpose (8), and the resulting masks were imported into Fiji (ImageJ) for fluorescence quantification. The mean fluorescence intensity of each cell was determined by dividing the raw integrated fluorescence density by the corresponding cell area. Background fluorescence was measured form cell-free regions of each image and subtracted from the mean fluorescence intensity of individual cells to obtain background-corrected fluorescence values.

**Bioinformatic Analysis.** Secondary structures of the full length *gnd/gndA* transcript were predicted using the RNAfold program (ViennaRNA 2.7.0) (9) and a customized version of the Afold software (10). The free energies of folding-unfolding ($\Delta G$) states for 100 nucleotide segments of RNA were estimated by Afold by calculating the difference between the optimal free energies of the paired (folded) and completely unpaired states of a given segment. The program was used to scan and process the entire *gnd/gndA* transcript sequence in all overlapping 100-nt windows. Temperature- and transcript-length-dependent changes in ensembles of suboptimal RNA structures were analyzed using the Hybrid program (11). Analyses were performed for the full-length *gnd/gndA* transcript, as well as for a series of truncated transcript sequences of increasing lengths, beginning at the *gnd* TSS (the first 100-nt window) and extending toward the transcript's 3' end in 100-nt window increments. A search for orthologous hits of the *gnd/gndA* transcripts was performed using the Basic Local Alignment Search Tool (BLAST; (12)) with the 'blastn' command (nucleotide BLAST+ v.2.6.0) against the NCBI 'prok_complete_genomes' and Prok2311 databases. Default parameters were used unless otherwise specified. Multiple nucleotide alignments of orthologous coding sequences were constructed using MUSCLE (13)and refined through pairwise comparisons using the OWEN program (14, 15). The resulting MSAs were folded using RNAalifold (ViennaRNA 2.7.0; (9)) to predict consensus RNA secondary structures. Unless otherwise specified, default parameters were used for RNA folding nd structure predictions; and minimum free energy (MFE) of predicted duplexes were also calculated. Infernal (v.1.1.2; (16)) was used for RNA sequence and structure analysis based on covariance models. Molecular phylogenetic analysis was performed using MEGA11 ((17); see Figure legends for details).

1. Yu D*, et al.* (2000) An efficient recombination system for chromosome engineering in *Escherichia coli*. *Proc Natl Acad Sci USA* 97:5978–5983.
2. Klinkert B*, et al.* (2012) Thermogenetic tools to monitor temperature-dependent gene expression in bacteria. *J Biotechnol* 160:55–63.
3. Melamed S, Adams PP, Zhang A, Zhang H, Storz G (2020) RNA-RNA interactomes of ProQ and Hfq reveal overlapping and competing roles. *Mol Cell* 77:411–425.e417.
4. Glaubig LC, Waldminghaus T, Naberhaus F (2011) Multiple layers of control govern expression of the *Escherichia coli ibpAB* heat-shock operon. *Microbiology (Reading)* 157:66–76.
5. Pienkoß S*, et al.* (2022) RNA thermometer-coordinated assembly of the *Yersinia* injectisome. *J Mol Biol* 434:167667.
6. Tan IS, Weiss CA, Popham DL, Ramamurthi KS (2015) A quality-vontrol mechanism removes unfit cells from a population of sporulating bacteria. *Dev Cell* 34:682–693.
7. Ng CY, Farasat I, Maranas CD, Salis HM (2015) Rational design of a synthetic Entner-Doudoroff pathway for improved and controllable NADPH regeneration. *Metab Eng* 29:86–96.
8. Pachitariu M, Rariden M, Stringer C (2025) Cellpose-SAM: superhuman generalization for cellular segmentation. *bioRxiv*.
9. Lorenz R*, et al.* (2011) ViennaRNA Package 2.0. *Algorithms Mol Biol* 6.
10. Ogurtsov AY, Shabalina SA, Kondrashov AS, Roytberg MA (2006) Analysis of internal loops within the RNA secondary structure in almost quadratic time. . *Bioinformatics* 22:1317–1324.
11. Matveeva OV, Shabalina SA (1993) Intermolecular mRNA-rRNA hybridization and the distribution of potential interaction regions in murine 18S rRNA. . *Nucleic Acids Res* 21:1007–1011.
12. Altschul SF, Gish W, Miller W, Myers EW, Lipman DJ (1990) Basic local alignment search tool. *J Mol Biol* 215:403–410.
13. Edgar RC (2022) High-accuracy alignment ensembles enable unbiased assessments of sequence homology and phylogeny. *Nat Commun13* 13:6968.
14. Ogurtsov AY, Roytberg MA, Shabalina SA, Kondrashov AS (2002) OWEN: aligning long collinear regions of genomes. *Bioinformatics* 18:1703–1704.
15. Kondrashov AS, Shabalina SA (2002) Classification of common conserved sequences in mammalian intergenic regions. . *Hum Mol Genet* 11:669–674.
16. Nawrocki E, Eddy SR (2013) Infernal 1.1: 100-fold faster RNA homology searches. *Bioinformatics* 29:2933–2935.
17. Tamura K, Stecher G, Kumar S (2021) MEGA11: Molecular Evolutionary Genetics Analysis Version 11. *Mol Biol Evol* 38:3022–3027.
18. Weaver J, Mohammad F, Buskirk AR, Storz G (2019) Identifying small proteins by ribosome profiling with stalled initiation complexes. *mBio* 10:e02819–02818.
19. Jones DT, Taylor WR, Thornton JM (1992) The rapid generation of mutation data matrices from protein sequences. *Comput Appl Biosci* 8:275–282.
20. Abramson J*, et al.* (2024) Accurate structure prediction of biomolecular interactions with AlphaFold 3. *Nature* 630:493–500.
21. Washietl S*, et al.* (2011) RNAcode: robust discrimination of coding and noncoding regions in comparative sequence data. *RNA* 17:578–594.

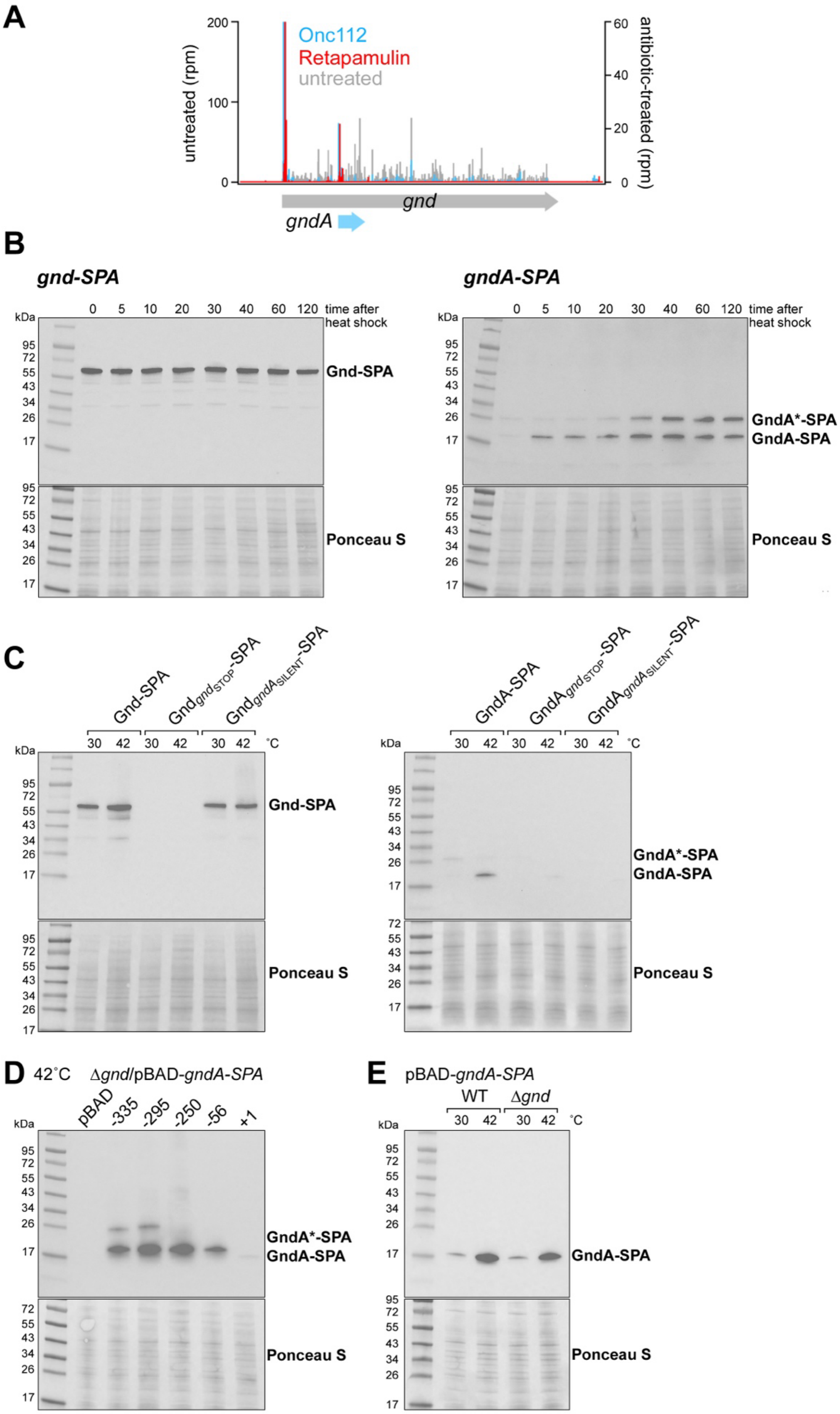
A
200
100
0
untreated (rpm)
Onc112
Retapamulin
untreated
60
40
20
0
antibiotic-treated (rpm)
gnd
gndA
B
gnd-SPA
gndA-SPA
kDa
0 5 10 20 30 40 60 120 time after heat shock
95 72 55 43 34 26 17
Gnd-SPA
GndA*-SPA
GndA-SPA
Ponceau S
C
Gnd-SPA
Gnd$_{gndSTOP}$-SPA
Gnd$_{gndASILENT}$-SPA
GndA-SPA
GndA$_{gndSTOP}$-SPA
GndA$_{gndASILENT}$-SPA
30 42 30 42 30 42 °C
D
42°C
Δgnd/pBAD-gndA-SPA
pBAD -335 -295 -250 -56 +1
E
pBAD-gndA-SPA
WT
Δgnd
30 42 30 42 °C

**Fig. S1.** (***A***) Browser images of ribosome profiling data for the *gnd* locus. Ribosome density for an untreated control (gray) and for cells treated with translation initiation-trapping antibiotics Onc112 (blue) or retapamulin (red) are shown. Data were taken from (18). (***B***) Western blot of Gnd-SPA or GndA-SPA levels after heat shock. Cells were grown in LB medium at 30˚C to mid-exponential phase ($OD_{600}$~0.5) and transferred to pre-warmed flask for heat shock at 42˚C. Samples were collected at the indicated time points. (***C***) Effects of *gndA* and *gndA* mutagenesis on protein levels. The $gnd_{STOP}$ mutation is a ATG-to-TGA substitution at the *gnd* start codon, whereas the $gndA_{SILENT}$ contains synonymous substitutions at codons D81, I98, and D99 that eliminate potential alternative start codons for *gndA*, without altering the Gnd amino acid sequence. Cells were grown in LB medium to mid-exponential phase ($OD_{600}$~0.5) at 30˚C, after which cultures were split with one half remained at 30˚C, while the other was subjected to heat shock at 42˚C for 20 min. (***D***) and (***E***) Western blot analysis of GndA-SPA variants with indicated N-terminal extensions. Constructs were expressed from the pBAD promoter in *E. coli* cells grown in LB medium at 30˚C to early exponential phase ($OD_{600}$ ~0.4). Expression was induced with 0.2% arabinose for 30 min. Cultures were then either transferred to a pre-warmed flask for heat shock at 42˚C for 20 min (D), or split into two aliquots, with one half kept at 30˚C and the other transferred to a pre-warmed flask at 42˚C for 20 min (E).

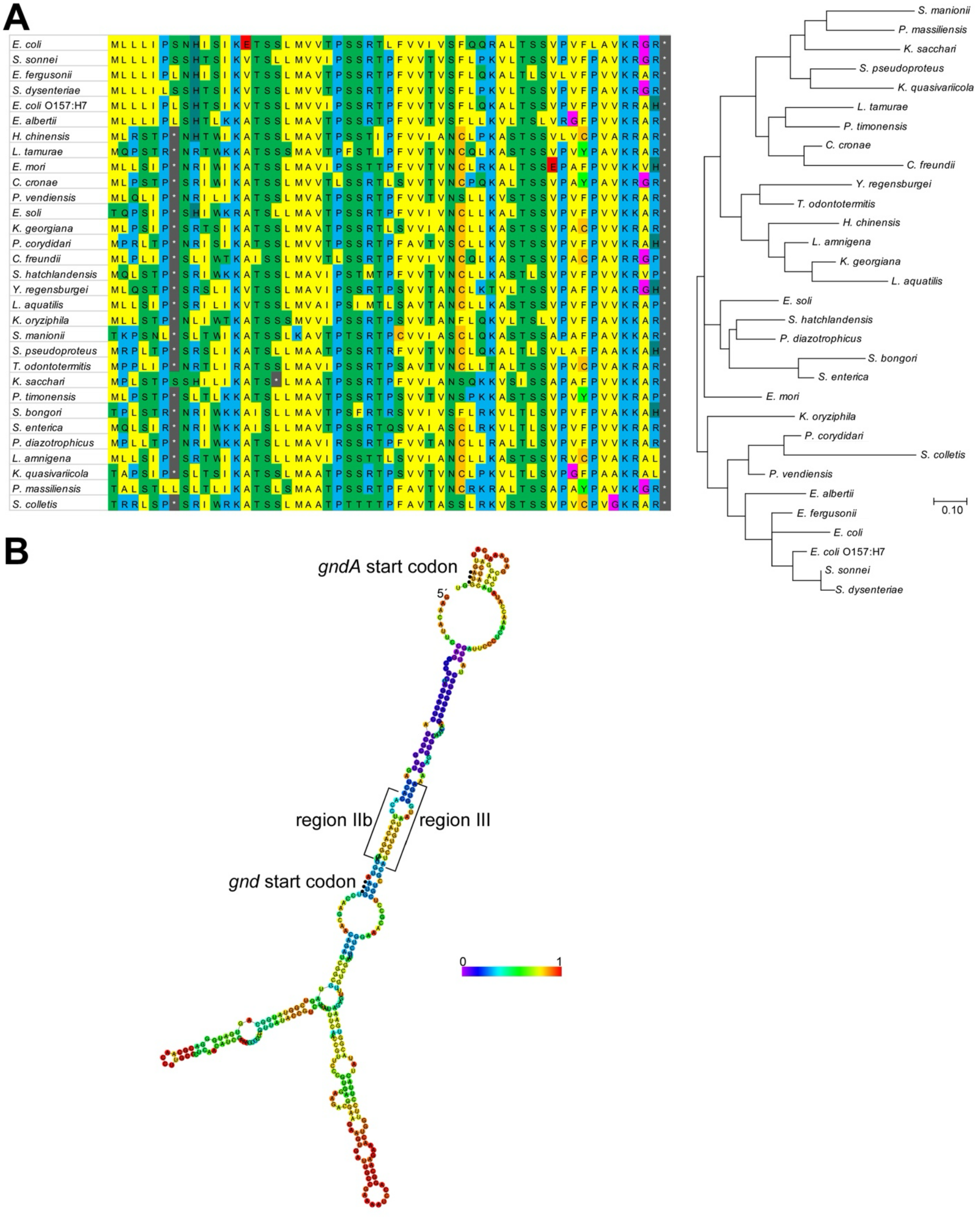


**Fig. S2.** Evolutionary analysis and phylogenetic trees of GndA orthologs. (***A***) Multiple sequence alignments (MSAs) of orthologous protein sequences were generated by translating the corresponding coding sequences and aligning them with MUSCLE. The resulting amino acid alignments were used to reconstruct evolutionary relationships and infer phylogenetic trees. Evolutionary analysis was inferred by using the Maximum Likelihood method and JTT matrix-based model (19). The tree with the highest log likelihood (-1719.26) is shown. Initial tree(s) for the heuristic search were obtained automatically by applying Neighbor-Join and BioNJ algorithms to a matrix of pairwise distances estimated using the JTT model and then selecting the topology with superior log likelihood value. A discrete Gamma distribution was used to model evolutionary rate differences among sites (5 categories (+G, parameter = 1.9405)). The tree is drawn to scale, with branch lengths measured in the number of substitutions per site. This analysis involved 32 amino acid sequences. The coding data was translated assuming a Standard genetic code table. All positions with less than 100% site coverage were eliminated, i.e., fewer than 0% alignment gaps, missing data, and ambiguous bases were allowed at any position (partial deletion option). There was a total of 51 positions in the final dataset. Evolutionary analyses were conducted in MEGA11 (17). The sequences used in the alignment and the tree are the following: *Escherichia coli* str. K-12 substr. MG1655 (NC 000913.3), *Shigella sonnei* (NZ CP055292.1), *Escherichia fergusonii* (NZ CP083638.1), Shigella dysenteriae (NZ CP055055.1), *Escherichia coli* O157:H7 str. Sakai (NC 002695.2), *Escherichia albertii* (NZ CP070290.1), *Huaxiibacter chinensis* (NZ JAJVHF010000021.1), *Leclercia tamurae* (NZ JAMHKR010000091.1), *Enterobacter mori* (NZ CP091779.1), *Citrobacter cronae* (NZ CP069763.1), *Pseudocitrobacter vendiensis* (NZ CALSBS010000005.1), *Enterobacter soli* (NC 015968.1), *Kluyvera georgiana* (NZ LSME01000020.1), *Pseudocitrobacter corydidari* (NZ CP087880.1), *Citrobacter freundii* (NZ CP033744.1), *Silvania hatchlandensis* (NZ JAMGZK010000046.1), *Yokenella regensburgei* (NZ RBIZ01000003.1), *Lelliottia aquatilis* (NZ JAENMQ010000001.1), *Kosakonia oryziphila* (NZ FMBC01000005.1), *Scandinavium manionii* (NZ JALIGC010000039.1), *Shimwellia pseudoproteus* (NZ WBXP01000081.1), *Trabulsiella odontotermitis* (NZ LIFU01000010.1), *Kosakonia sacchari* (NZ CP016337.1), *Leclercia pneumoniae* (NZ CP071383.1), *Pseudenterobacter timonensis* (NZ FCOP01000013.1), *Salmonella bongori* N268-08 (NC 021870.1), *Salmonella enterica* subsp. *enterica* serovar Typhimurium str. LT2 (NC 003197.2*),* *Phytobacter diazotrophicus* (NZ CP051548.1), *Lelliottia amnigena* (NZ CP079896.1), *Klebsiella quasivariicola* (NZ CP084768.1), *Phytobacter massiliensi*s (NZ CABKSF010000001.1), and *Siccibacter colletis* (NZ CP074352.1). ***(B)*** RNA folding predictions (9) for the first 356 nt of the *gnd* transcript is consistent with a highly stable interaction between the *gnd* ribosome binding site and start codon (region IIb) and the ICS (region III) and relaxed secondary structure in the vicinity of the *gndA* ribosome binding site and start codon. Start codons of *gnd* and *gndA* ORFs are marked by blue circles. Coloring scheme shows base-pair probabilities from 0 to 1.

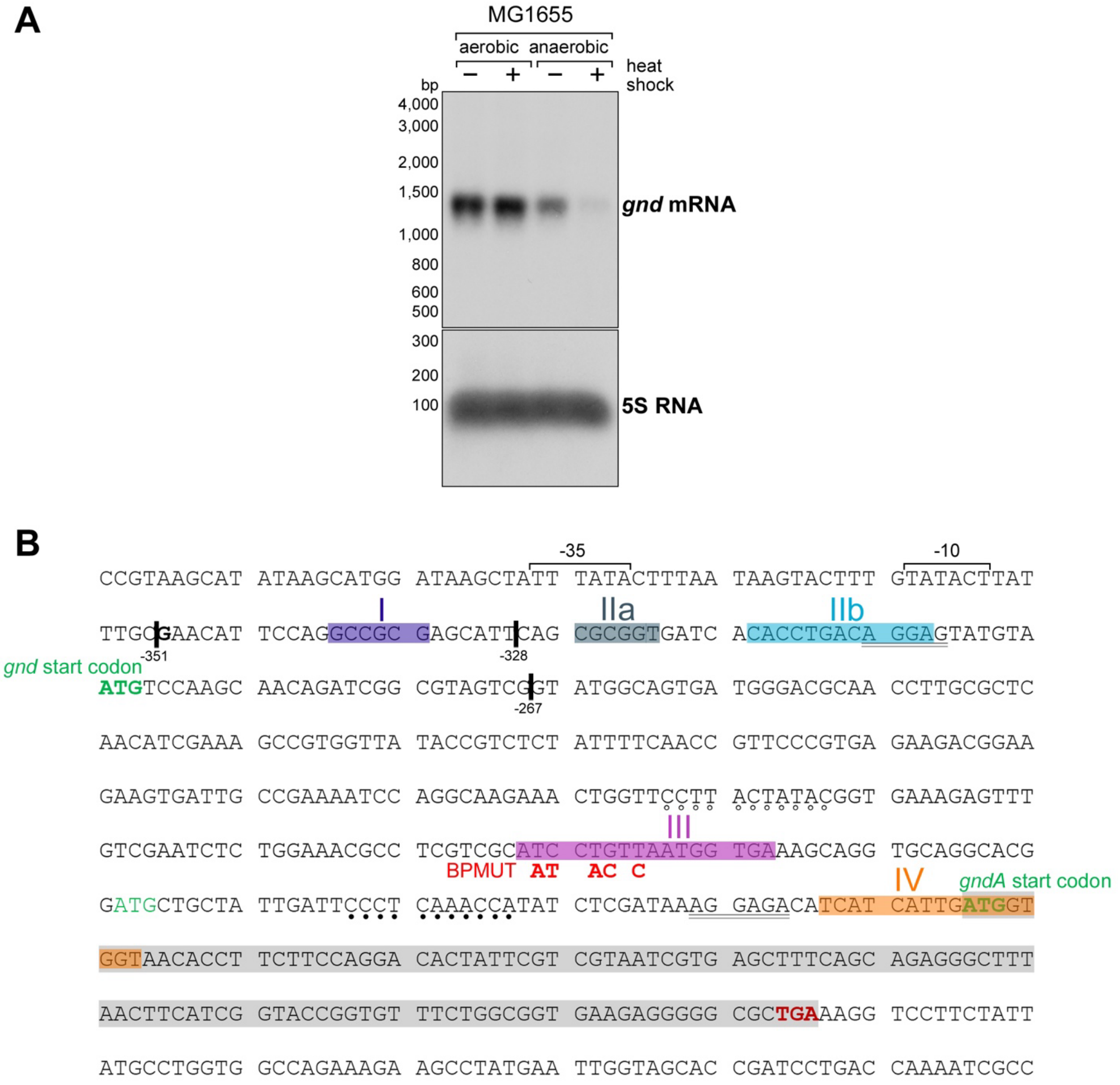


**Fig. S3.** Levels and sequence of *gnd* mRNA. **(*A*)** Northern analysis using an oligonucleotide probe specific to the *gnd* locus. Total RNA was isolated from cells grown in LB medium to mid-exponential phase ($OD_{600}$~0.5) at 30˚C under either aerobic or anaerobic conditions. Cultures were then divided into two aliquots; one remained at 30˚C, while the other was subjected to heat shock at 42˚C for 20 min. RNA samples were subsequently analyzed by Northern blot. **(*B*)** Coding sequence of *gndA* and 300 nt upstream. Coloring for Gnd and $GndA_{36}$-SPA codons is as in Fig. 1A. Region 1, IIa, IIb, III, and IV sequences involved in base pairing are highlighted in different colors. Black lines indicate 5´ ends of fusions assayed in **Fig. 2C**. Mutations to disrupt pairing with region III are in red font. Sequences listed in Fig. 2D are indicated with black dots (increased modification) and white dots (decreased modification).

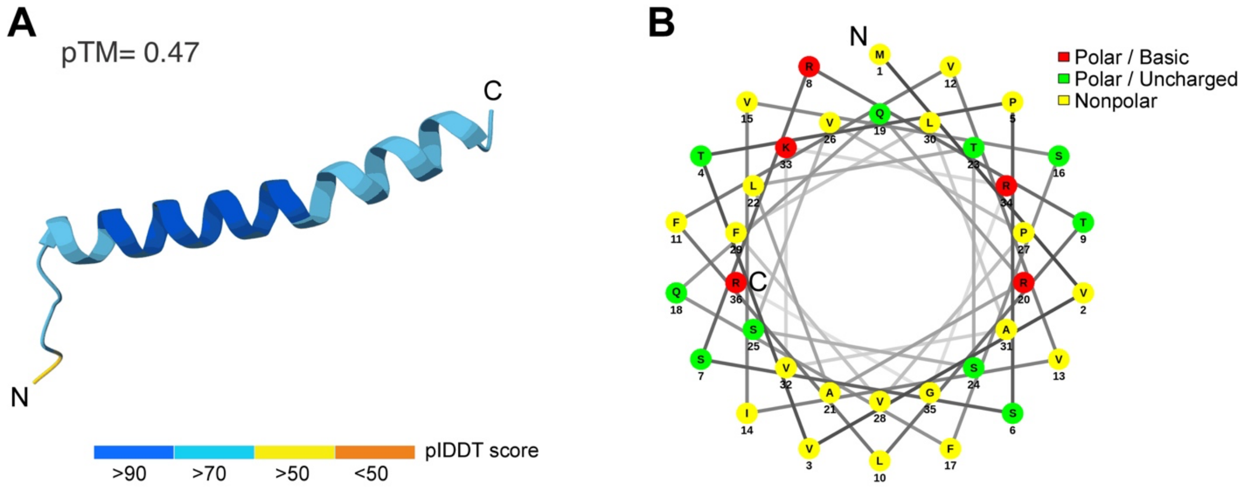


**Fig. S4. (*A*)** Predicted structure of GndA generated by AlphaFold3 (20) on February 16, 2025. The model is colored according to the predicted local distance difference test (pIDDT) confidence score, as indicated. The predicted TM-score (pTM) is shown above the model. **(*B*)** Helical wheel projection of the predicted α-helix, illustrating the distribution of amino acid side chains around the helix. Residues are colored according to their physicochemical properties: polar/basic (red), polar/uncharged (green), and nonpolar (yellow). The clustering of hydrophobic residues on one face of the helix and polar residues on the opposite face predicts that GndA forms an amphipathic α-helix.

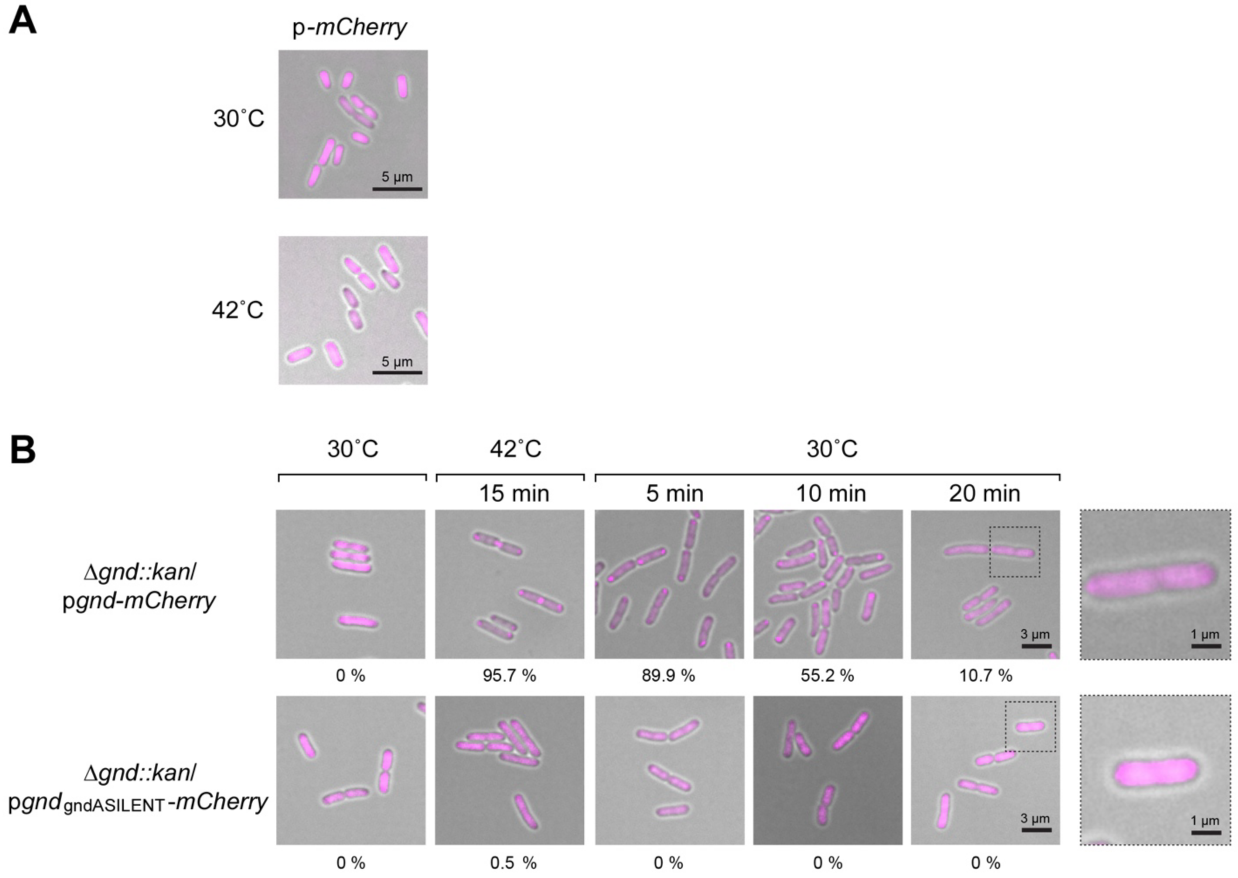


**Fig. S5. (*A*)** Expression of plasmid-borne mCherry. Representative epifluorescence microscopy images of cells expressing mCherry from the T7 promoter. Cells were grown in LB medium at 30ºC to mid exponential phase ($OD_{600}$~0.5). mCherry expression was an induced with 1 mM IPTG for 15 min before cultures were split into two aliquots, where one half maintained at 30˚C (upper panel) and the other exposed to heat shock at 42˚C for 5 min (lower panel) prior to imaging. **(*B*)** Representative fluorescence microscopy images of *E. coli* BL21 Δ*gnd* cells carrying a plasmid-borne *gnd-mCherry* fusion or $gnd_{gndASILENT}$-*mCherry* fusion showing increasing polar localization with time after heat shock and decreasing polar localization with time after return to 30˚C. Cells were grown in LB medium supplemented with glucose at 30˚C to early exponential phase ($OD_{600}$~ 0.35). Expression was induced with 0.5 mM IPTG for 15 min. Cells were then collected and washed to remove the inducer. One fraction was resuspended in 30˚C LB and a sample was imaged. The other fraction as resuspended in 42˚C LB, and samples were taken at 15 min. Subsequently, the culture was shifted back to 30˚C and samples were taken at 5, 10 and 20 min. Scale bar for left four columns = 3 μm. Polar localization for these samples was quantitated for >300 cells for each sample and the percent cells with polar localization is indicated below each image. Right column shows enlargement of cells in regions boxed by dotted line. Here scale bar = 1 μm.

| | *gndA* | *gndA* | *ftsO* | *ftsO* |
|---|---|---|---|---|
| **Methods** | **Score** | ***P-value*** | **Score** | ***P-value*** |
| RNAcode (high coding potential) * | 0.8 | 0.01 | 0.1 | 0.53 |
| Base-pair distance, consensus | -1.433 | 0.11 | -2.449 | 0.02 |
| RNAdistance, (coarse-grained), consensus | -0.535 | 0.31 | -2.269 | 0.02 |
| RNAdistance (HIT), consensus | -0.469 | 0.3 | -2.135 | 0.01 |
| RNAdistance, (w. coarse-grained), consensus | -0.372 | 0.37 | -2.084 | 0.04 |
| RNAdistance, (w. coarse-grained), consensus | -0.277 | 0.42 | -2.038 | 0.03 |
| Base-pair distance, pairwise | -0.219 | 0.33 | -2.029 | 0.02 |
| RNAdistance (coarse-grained), consensus | -0.182 | 0.45 | -2.021 | 0.02 |
| RNAdistance (full), consensus | -0.177 | 0.5 | -1.956 | 0.01 |
| RNAdistance (coarse-grained), pairwise | -0.162 | 0.39 | -1.862 | 0.02 |
| Ensemble distance, consensus | -0.064 | 0.56 | -1.739 | 0.06 |
| Ensemble distance, pairwise | -0.221 | 0.36 | -2.035 | 0.04 |
| RNApdist-like measure, pairwise | -0.154 | 0.38 | -1.748 | 0.05 |
| RNApdist-like measure, consensus | -0.163 | 0.5 | -1.951 | 0.01 |
| Mountain metric using base-pair probabilities, pairwise | 0.382 | 0.57 | -1.267 | 0.08 |
| Mountain metric using base-pair probabilities, consensus | -0.431 | 0.41 | -2.101 | 0.03 |
| SCI (structural selective index) | 1.452 | 0.07 | 2.053 | 0.01 |
| pairwise SCI using RNAeval | 1.912 | 0.05 | 2.647 | 0 |

**Table S1. Structure conservation analysis (SCA) and estimation of coding potential potential.** Different approaches were used to assess protein-coding potential (marked with *) and RNA structural selective constraints. Protein-coding potential was evaluated using RNAcode (21), whereas RNA structural constraints were assessed primarily based on RNA features using the Vienna package (9). Salmon fill highlights the pronounced differences in assigned scores between the GndA small protein and the FtsO small RNA, both of which were detected in annotated protein coding regions.
